# Cytocapsular tubes and networks function as physical superdefence freeway systems conducting conventional cancer drug pan-resistant tumor metastasis

**DOI:** 10.1101/2022.01.05.475158

**Authors:** Tingfang Yi, Gerhard Wagner

**Author notes:** Corresponding author: Dr. Tingfang Yi, Cytocapsula Research Institute, 245 First Street, Cambridge, MA, 02142 USA, or at.

## Abstract

Cancer drug pan-resistant tumor metastasis (cdp-rtm) is a major source of cancer lethality. Cytocapsular tubes (CCTs) and their networks are physical membrane-enclosed freeway systems for cancer cell dissemination across tissues and organs *in vivo*. Whether cytocapsular tube superlarge biomembranes function as superdenfence and conduct cdp-rtm is unknown. It is also unknown whether conventional cancer drug development methods, including cancer cell line derived xenograft (CDX) and patient cancer cell derived xenograft (PDX), generate cytocapsular tubes (CCTs). It is also unclear whether xenografts can be created that contain CCTs for efficient cancer drug development. Here, we investigated CCT functions related to cancer drug resistance, CCTs in CDX and PDX and CCT xenograft (CCTX). Using clinical cancer tissues, we discovered that CCTs potently shielded against multiple chemotherapy treatments with diverse conventional cancer drugs. Next, our quantitative analyses show that CCT biomembrane drug barriers significantly increase cancer drug resistance by 6.6-folds to14-folds. We found that conventional CDX and PDX animal models do not generate CCTs in these xenografts. By mimicking *in vivo* cancer cell environments for cancer patient cancer cell culturing, we have successfully isolated CH-5^high^/CH-6^high^ subpopulations of patient breast cancer cells and pancreas cancer cells that are propertied with cytocapsular tube generation capacities and engender large quantities of CCTs in mouse xenografts. Biochemical and immunohistochemistry analyses demonstrated that CCTs in these xenografts are similar to those in clinical cancer tissues. In summary, our research has identified that CCTs and networks function as physical superdefence freeway systems conducting conventional cancer drug pan-resistant tumor metastasis, and developed a CCTX platform for highly efficient cancer drug development, which pave avenues for more efficient development of effective and precise cancer drugs for tumor cure at both personal and broad-spectrum levels.

## Introduction

Cancer is a leading source of human lethality in the world. It is estimated that approximately 10million of cancer deaths and approximately 20million new cancer cases occur in 2020 alone worldwide^1^. The global cost of cancer was approximately $2.5 trillion in 2010 alone.^2^ There are approximately 5million published articles on solid and liquid cancers since 1783 (in PubMed). Furthermore, US FDA approved about 650 cancer drugs since1949, and there were centuries’ intensive efforts and innumerous spending; however, the marginal outcomes suggest the extreme complexity of cancer and that the major parts of the nature of cancer are still beyond our conventional understanding and knowledge^3–6^.

The integrated cancer metastasis and pan-drug resistance are major sources of cancer lethality. Most cancer patients with disseminated cancer die because these metastases become resistant to all available drugs, which leads to biological function failure of multiple tissues and organs. Frequently, cancer drug resistance arises in two steps: initially, the tumors variably respond to the cancer drug, but not all cancer cells are killed. Subsequently, the surviving cancer cells develop into new tumors that do not respond to any drug anymore^7–8^. Based on tumor response to the initial therapy, cancer resistance can be broadly classified into two categories of primary and acquired resistances: primary cancer drug resistance exists prior to treatment, while acquired resistance occurs after initial therapy^9^. Multiple mechanisms of cancer drug resistances are reported and intensively investigated, including: modification of drug transports, mutation of extracellular receptors, amplification and mutation of drug molecular targets, drug metabolism (uptake, efflux and detoxification), mutation of cancer cell addicted genes/proteins, amplification of alternative oncogenes, activation of alternative survival pathways, multidrug resistance, cross resistance (resistance to one drug leads to resistance to other drugs), and so on^10–14^. Multiple hypotheses on mechanisms of cancer metastasis had been raised, including: seed-soil theory^4^, metastatic cascade theory (with five steps of invasion, intravasation, circulation, extravasation and colonization)^3,15^, cancer stem cell (CSC) theory^4^, and so on. In these hypotheses, metastatic cancer cells must overcome numerous physical obstacles barring metastasis and must invade vasculature to reach far distance destinations, which are controversial to clinical observations: there are no circulating tumor cells in the blood in solid cancer stage I and II in which millions or hundreds of millions of cancer cells have already metastasized to far distance sites diagnosed as revealed with novel precise cancer diagnosis methods^16–18^.

Recently, we discovered that single cancer cells can generate superlarge, superlong (up to 110m in length), membrane-enclosed tube-shaped cytocapsular tubes (CCTs) outside of the cell, and provide physical highways free of ECM obstacles for highly efficient cancer cell dissemination across tissue and organs^17^. Furthermore, we discovered that CCTs are universally present in all 202 kinds (types and subtypes) of checked solid cancers, but not in normal tissues or benign tumors^17^. We found that cancer metastasis occurs at the initiation stage, much earlier than the conventional cancer diagnosis. Even at the breast cancer *in-situ* stage (conventionally thought the initiation of cancer before invasion), there are thousands of CCTs and breast cancer cells that already aggressively migrate and invade in CCTs in local tissues. We found that CCTs and its networks, other than circulation systems, dominate cancer metastasis pathways in all of the investigated 202 kinds (including subtypes) of clinical cancers^17^. Only at the late stages (after Stage IIc), occasionally, a few CCTs invade into micro blood vessels and lymph vessels and release cancer cells or fragments of cancer cells into the circulation vessels, which are the resource for circulating tumor cell DNA detection. It is estimated that only about 1 in 100million cancer cells is released into circulation systems after Stage IIc^16–18^. During cancer pharmacotherapy, drug resistant cancer metastasis leads to cancer lethality. It is unknown whether CCTs and its networks function as drug barriers to conventional cancer drug panresistance.

The efficient development of effective and precise cancer drugs depends on whether cancer drugs can overcome drug-resistant tumor metastasis as provided by CCTs. It is unknown whether CCTs are present in the popularly employed cancer drug screening tools of 2D cell and 3D cell colonies and in organoids *in vitro*, and in CDX and PDX in animal models *in vivo.* Here, we have comprehensively investigated CCT functions in conventional cancer drug resistance, asked whether conventional CDX and PDX generate CCTs, and methods to invent xenografts with CCTs. We found that CCT membranes provide a superdenfence mechanism for disseminating cancer cells and display pan-resistance to conventional cancer drugs. In contrast, conventional CDX and PDX animal models in mouse do not generate CCTs. Thus, we created a CCTX mouse model to engender high density CCTs and networks similar to those found in clinical cancer tissues. Our studies pave avenues for the efficient development of effective and precise cancer drugs for the cure of cancers at both the personal and broad-spectrum levels.

## Results

### The membranes of cytocapsular tubes and their networks function as superdefence drug barriers and create freeway systems that permit cancer cell metastasis resistant to conventional cancer drugs

Metastasis of drug pan-resistant tumor cells is a major source of cancer lethality^2^, and cytocapsular tubes (CCTs) provide the physical pathways for cancer cell metasis^16–18^. In order to investigate whether the bi-layer phospholipid bio-membrane composed superlarge CCT membranes function as physical barriers to cancer drugs, we examined FFPE (formalin-fixed paraffin-embedded) cancer tissue specimens from a Stage IIc breast cancer patient who experienced 3 chemotherapies with 3 cancer drugs of Pertuzumab, Palbociclib, and Docetaxel. CT-scan diagnosis showed that the biopsy area is clear, 1 month after the 3^rd^ chemotherapy. However, with CCT immunohistochemistry fluorescence microscope analyses, we observed that: 1) there are a lot of curved and winded CCTs which pile up and form compact CCT masses, 2) cancer cells are in migration in CCTs, 3) there are no cancer cell masses located outside of CCTs, and 4) there are many big CCT cavities (**Fig. 1A**). These observations evidenced that: 1) cancer drugs induce degradation of parts of CCTs and cause creation of many big CCT cavities; 2) some CCTs respond to these chemotherapies, but the major parts of CCTs remain intact, 3) the remaining CCTs display pan-resistance against conventional cancer drugs, 4) breast cancer cells invade into CCTs and superlarge CCT membranes protect cancer cells by retaining cancer drugs outside of CCTs.

**Fig.1.**
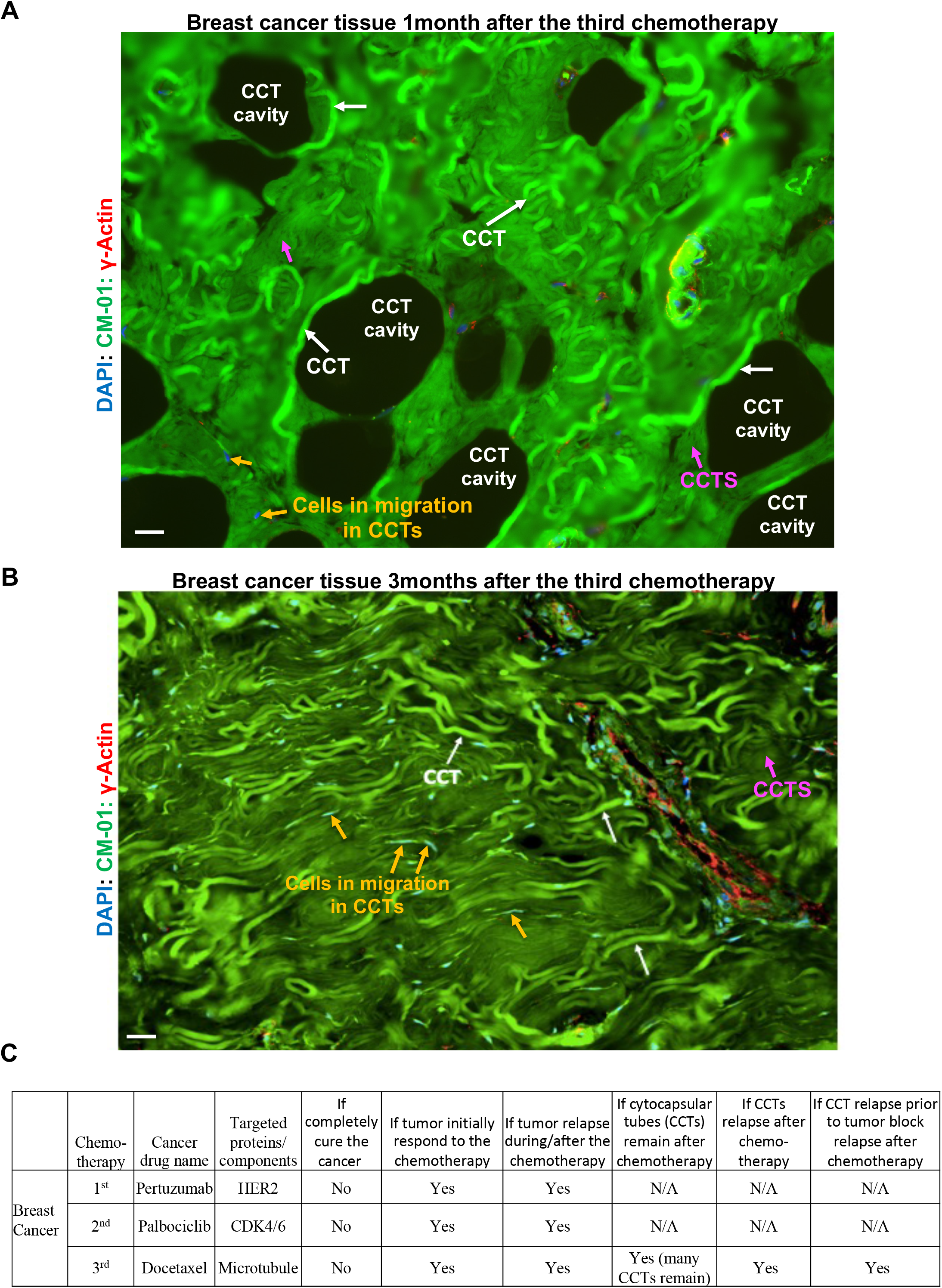
Cytocapsular tubes and networks function as both cancer metastasis physical pathways and superdefence of cancer cells inducing cancer drug pan-resistance in clinical breast cancer. (A) Representative fluorescence microscope image of needle biopsy breast cancer tissue 1 month after the third chemotherapy. Three cycles of chemotherapies were performed with three cancer drugs of Pertuzumab, Palbociclib and Docertaxel, respectively. The FFPE tissue slides were performed with immunohistochemistry fluorescence staining with anti-CM-01 and anti-γ-actin antibodies and DAPI. A lot of curved and winded cytocapsular tubes (CCTs, white arrows) pile together and form compact CCT masses, in which there are many big CCT cavities caused by CCT degradation, decomposition and disappearance. Cancer cells in migration in CCTs (orange arrows) and CCT strands (CCTS, purple arrows) are shown. Scale bar, 10μm. (B) Representative fluorescence microscope image of needle biopsy breast cancer tissue 3 months after the third chemotherapy (from the same female patient as in A). There are a lot of curved and winded and non-degraded cytocapsular tubes (CCTs, white arrows) pile together and form compact CCT masses, and fill all CCT cavities with no CCT cavity left. Cancer cell in migration in CCTs (orange arrows) and CCT strands (CCTS, purple arrows) are shown. Scale bar, 10μm. (C) Table of the CCT analyses after the three chemotherapies. (Note: N/A, not available; these analyses are not performed and the results are not available).

Next, we further investigated the breast cancer tissue specimens of the same patient taken 3 months after the 3^rd^ chemotherapy. We found that: 1) there are have much more CCTs and all previous CCT cavities are filled or covered by dense CCTs, 2) most (>95%) of the CCTs appear in regular CCT early-stage without degradation, 3) more breast cancer cells are in migration in the large quantities of CCTs, 4) most cancer cells are in CCTs but not located outside of CCTs (**Fig.1B**). Combined together (**Fig. 1A-1C**), these observations demonstrated that CCT superlarge membranes function as physical superdefence and freeway systems conducting conventional CDP-RTM.

Next, we investigated whether CCTs are in volved in cdp-rtm caused cancer lethality. We examined CCTs in the FFPE autopsy specimens of metastatic prostate cancers in the bladder from a dead prostate cancer patient who previously received several chemotherapies, and the last three chemotherapy drugs were Enzalutamide, ZYTIGA™, and Cabazitaxel. We observed that: 1) there are many metastatic prostate cancer CCTs in the bladder, 2) most CCTs are at late stage and in degradation, 3) bladder tissues lost the normal tissue texture and cell density (**Fig. 2A-B**). These observations evidenced that: 1) prostate cancer-cell CCTs and networks provide with cancer drug pan-resistant metastatic physical pathways for prostate cancer cells to disseminate into the bladder, 2) metastatic prostate CCTs and prostate cancer cell invaded into the bladder leading to failure of the bladder biological function, 3) prostate cancer cell via CCT drug panresistant metastatic freeways invade into multiple tissues/organs, which leads to: new tumor growth, failure of biological function of multiple organs, and subsequent patient death.

**Fig.2.**
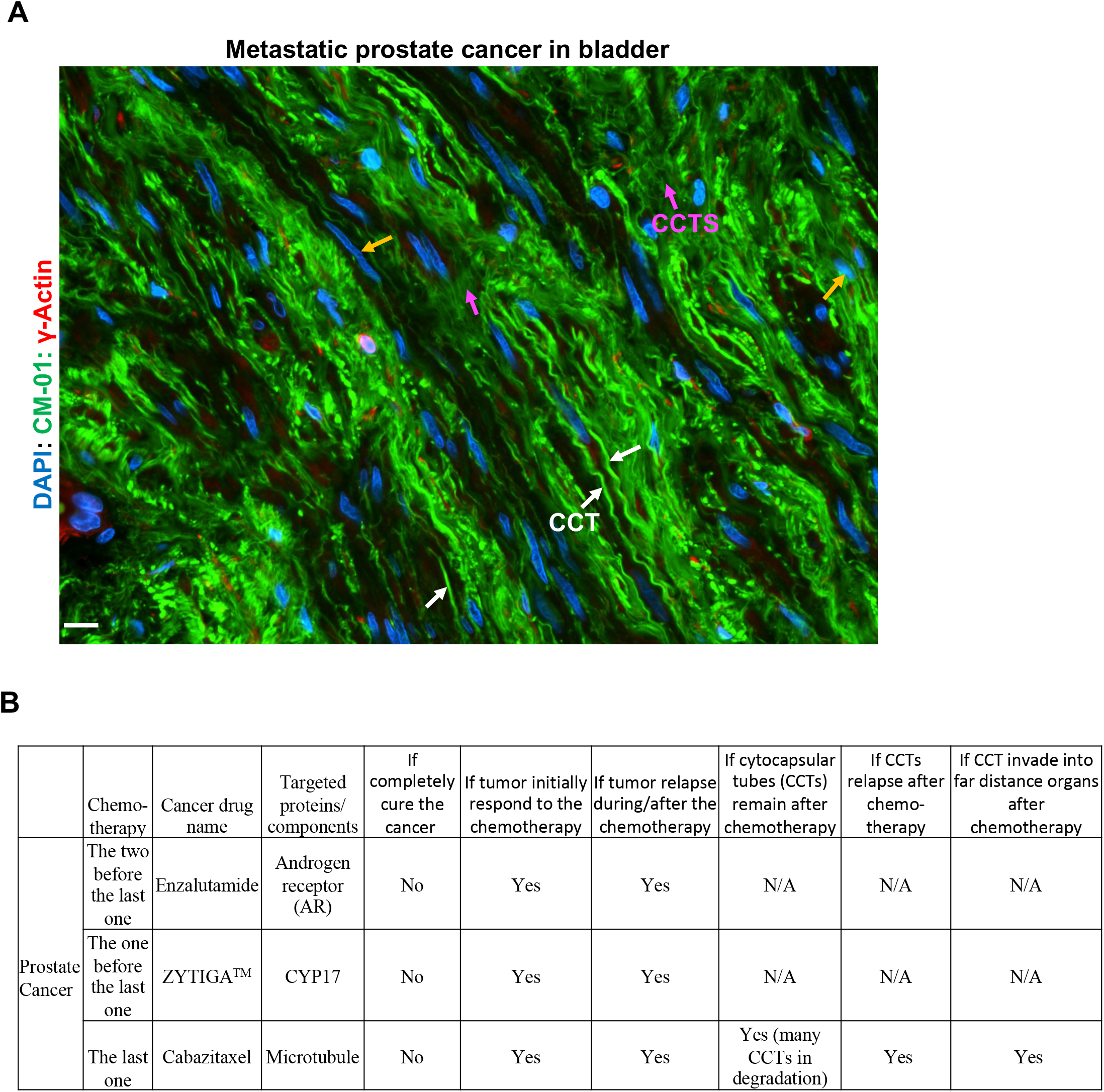
Cytocapsular tubes and networks function as both cancer metastasis physical pathways and superdefence of cancer cells inducing cancer drug pan-resistance in clinical prostate cancer. (A) Representative fluorescence microscope image of autopsy metastatic prostate cancer tissues after several chemotherapies. The last three chemotherapies were performed with three cancer drugs of Enzalutamide, ZYTIGA™, Cabazitaxel, respectively. The FFPE tissue slides were performed with immunohistochemistry fluorescence staining with anti-CM-01 and anti-γ-actin antibodies and DAPI. A lot of cytocapsular tubes (CCTs, white arrows) are in degradation and degrade into thick, thin or very thin CCT strands. Cancer cells in migration in CCTs (orange arrows) and CCT strands (CCTS, purple arrows) are shown. Scale bar, 10μm. (B) Table of the CCT analyses after the last three chemotherapies. (Note: N/A, not available; these analyses are not performed and the results are not available).

In summary, these observations (**Fig. 1-2**) and the previous data of 7,125 cancer patients (including 6,928 dead cancer patients)^17^ demonstrated that cancer cell CCTs and their networks function as physical superdefence mechanisms and provide freeways to conduct metastasis of tumors that are pan-resistant to conventional cancer drugs. They lead to cancer metastasis to multiple organs and subsequent biological function failure, and cause cancer lethality. Most commercial conventional cancer drugs do not successfully inhibit CCT conducted cancer metastasis, and fail to save cancer patients’ lives. These data show that the conventional low efficient cancer drug development methods may miss the superlarge CCTs and their networks which function as both superdefence and tumor metastasis freeways, in the preclinical screening procedures.

### Superlarge and enclosed cytocapsular tube membranes shield cancer drugs outside and significantly increased cancer cells drug resistance

In order to further quantify the CCT barrier effects on cancer drugs, we investigated the cancer drug barrier effects of cancer cell CCT membranes of multiple cancer cells with multiple commercial cancer drugs. The cell viability IC_50_ of Sunitinib (targeting multiple receptor tyrosine kinases, RTKs) on 2D pancreas cancer Bxpc3 cells without CCTs is 4.82±0.6μM, while on Bxpc3 cells with and in CCTs (on 3D Matrigel matrix) is 31.68±2.09μM, increasing 6-folds (**Fig. 3A-C**). The cell viability IC_50_ of Apalutamide (targeting Androgen receptor, AR) on 2D prostate cancer LnCap cells without CCTs is 0.51±0.03μM, while on LnCap cells with and in CCTs is 4.2±0.12μM, increasing 8-folds. The cell viability IC_50_ of Abiraterone acetate (targeting CYP17A1) on 2D prostate cancer LnCap cells without CCTs is 1.17±0.63μM, while on LnCap cells with and in CCTs is 16.43±2.34μM, increasing 14-folds (**Fig. 3C**). These data evidenced that cancer cell CCTs substantially shield conventional cancer drugs outside, largely block conventional cancer drugs to access cancer cells in CCT lumens, and potently increase drug resistance of cancer cells.

**Fig.3.**
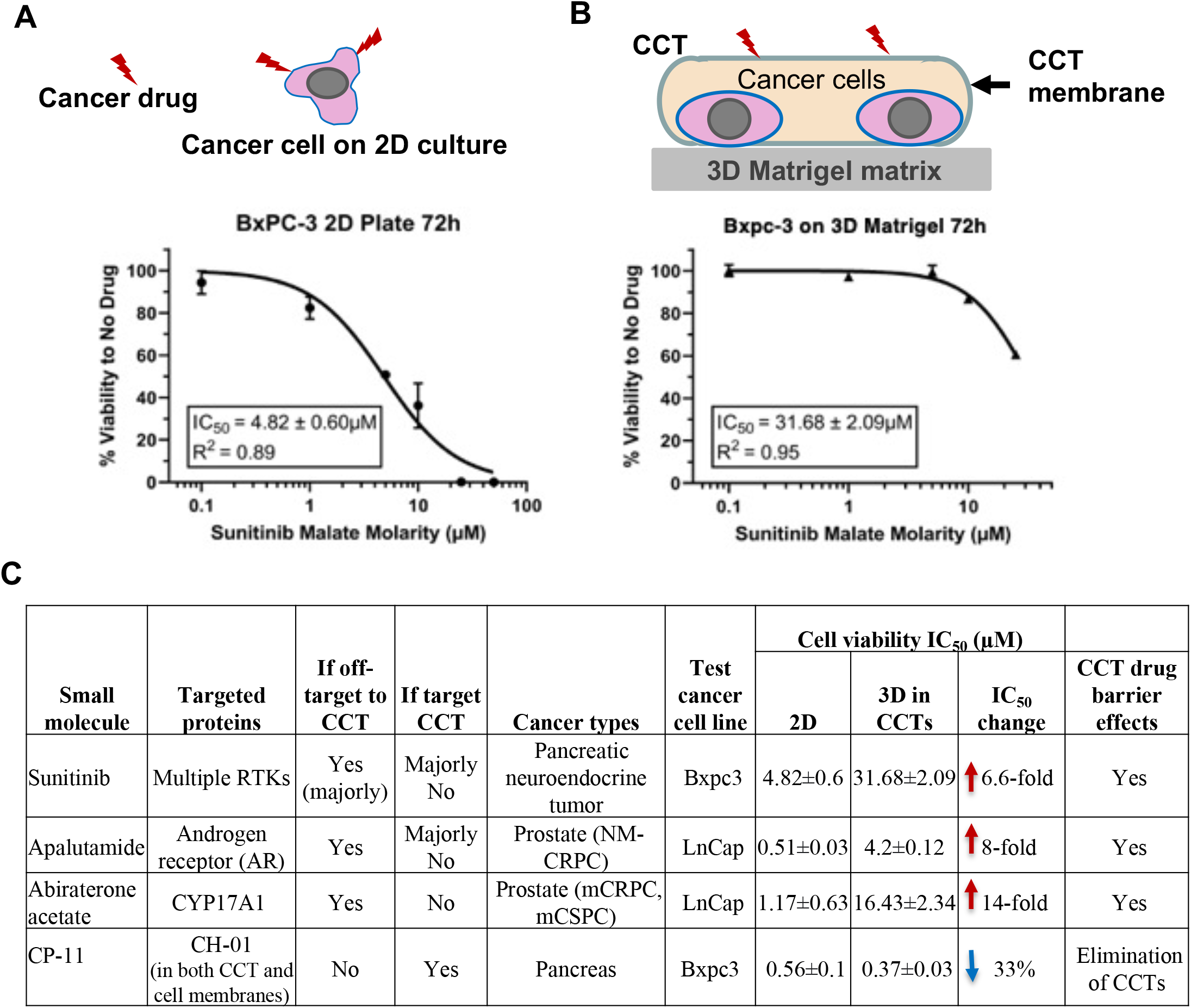
Quantitative analyses of cytocapsular tube superlarge membrane drug barrier effects to multiple cancer drugs and effects of small molecule CP-11 targeting CCT membrane protein CH-01 on CCT superdefence. (A) Graph of ATP-based cell viability assays showing the effects of small molecule of cancer drug Sunitinib on pancreas cancer cell Bxpc3 on 2D cell culturing. Top panel shows that small molecule cancer drugs directly access cancer cell cytoplasm membranes. (B) Graph of ATP-based cell viability assays showing the effects of small molecule of cancer drug Sunitinib on pancreas cancer cell Bxpc3 on 3D Matrigel matrix in which single Bxpc3 cancer cells generate cytocapsulae and CCTs. Top panel shows that the superlarge CCT bi-layer phospholipid membranes wrap Bxpc3 cancer cells inside, shield small molecule cancer drug Sunitinib outside, and decrease Sunitinib to access cancer cells in CCT lumens. (C) Table showing the effects of multiple small molecules cancer drugs of Sunitinib, Apalutamide, Abiraterone acetate, and CCT membrane protein CH-01 targeted small molecule CP-11 on the indicated cancer cells on 2D and 3D in CCTs. Triplicated ATP-based cell viability assays of each small molecule were performed. The average cell viability IC_50_ results are shown and compared.

Next, we investigated if small molecules that target cancer cell CCT important molecular markers can effectively inhibit CCT drug barrier effects. We developed a small molecule of CP-11 that binds and blocks the active center of CCT key molecular marker membrane protein CH-01 (which displays high abundance in CCT membranes of cancer cells in >150 kinds examined clinical solid cancers. Its relative abundance in normal cells, cancer cells and cancer cell CCTs is about 1:8:100). In contrast and not unexpected, the cell viability IC_50_ of CP-11 on 2D pancreas cancer Bxpc3 cells is 0.56±0.1μM, while on Bxpc3 cells with and in CCTs is 0.37±0.03μM, decreasing 33% (**Fig. 3C**). These data demonstrated that small molecules that target CCT key molecule protein markers can effectively eliminate CCT superdefence and kill cancer cells in CCT lumens.

In summary, these quantitative data (**Fig. 3**) evidenced that cancer cell CCTs and their networks substantially shield conventional cancer drugs outside, reducing access to cancer cells in CCTs. This effect significantly increased resistance to broad-spectrum conventional cancer cells drugs, and lead to conventional cancer drug pan-resistance. These results (**Fig. 3**) suggested that small molecules that target CCT key genes/proteins can potentially abolish CCT superdefence and eradicate cancer cells in CCTs and its networks in clinical chemotherapy.

### CCTs are absent in conventional cancer drug development methods CDX and PDX

Efficient *in vitro* screening methods of cancer drug candidates are essential for effective and efficient cancer drug development. Next, we investigated whether cancer cells in cancer cell line derived xenograft (CDX)^3–4^ and patient cancer derived xenograft (PDX)^3–7^ generate CCTs. In the CDX models (1million cancer cells/injection) of 3 breast cancer cells lines (MDA-MB-231, MCF-7 and BT-474) and the pancreas cancer cell line of Bxpc3, 100% of injections formed tumors. However, none of these CDX tumors developed CCTs (**Fig. 4A, 5A-B**). In the PDX models of breast cancer and pancreas cancer (1million cancer cell/injection/mouse, 5mice/group), the tumor formation percentages are only 40% and 20%, respectively. Consistently, none of these PDX tumors show CCTs (**Fig. 4B, 5C-D**). In the PDX models of breast cancer and pancreas cancer (2million cancer cell/injection/mouse, 5mice/group), the tumor formation percentages increased to 100% and 80%, respectively. Constantly, none of these PDX tumors generate CCTs (**Fig. 4B, 5C-D**). These data evidenced that the conventional CDX and PDX tumors in animal models *in vivo* tested here, do not generate CCTs. This demonstrated that the cancer drug candidates, developed through these conventional screening models in which CCTs are absent, will not have the capacity by design to eliminate CCT superdefence and eradiate cancer cells in CCTs and their networks. These data are consistent with the current facts: 1) the very low (average only 3%) successful rate for prediction of clinical value based on cell lines and animal models without CCTs in development of current cancer drugs; and 2) the weak chemotherapy outcomes in clinical cancer therapy.

**Fig.4.**
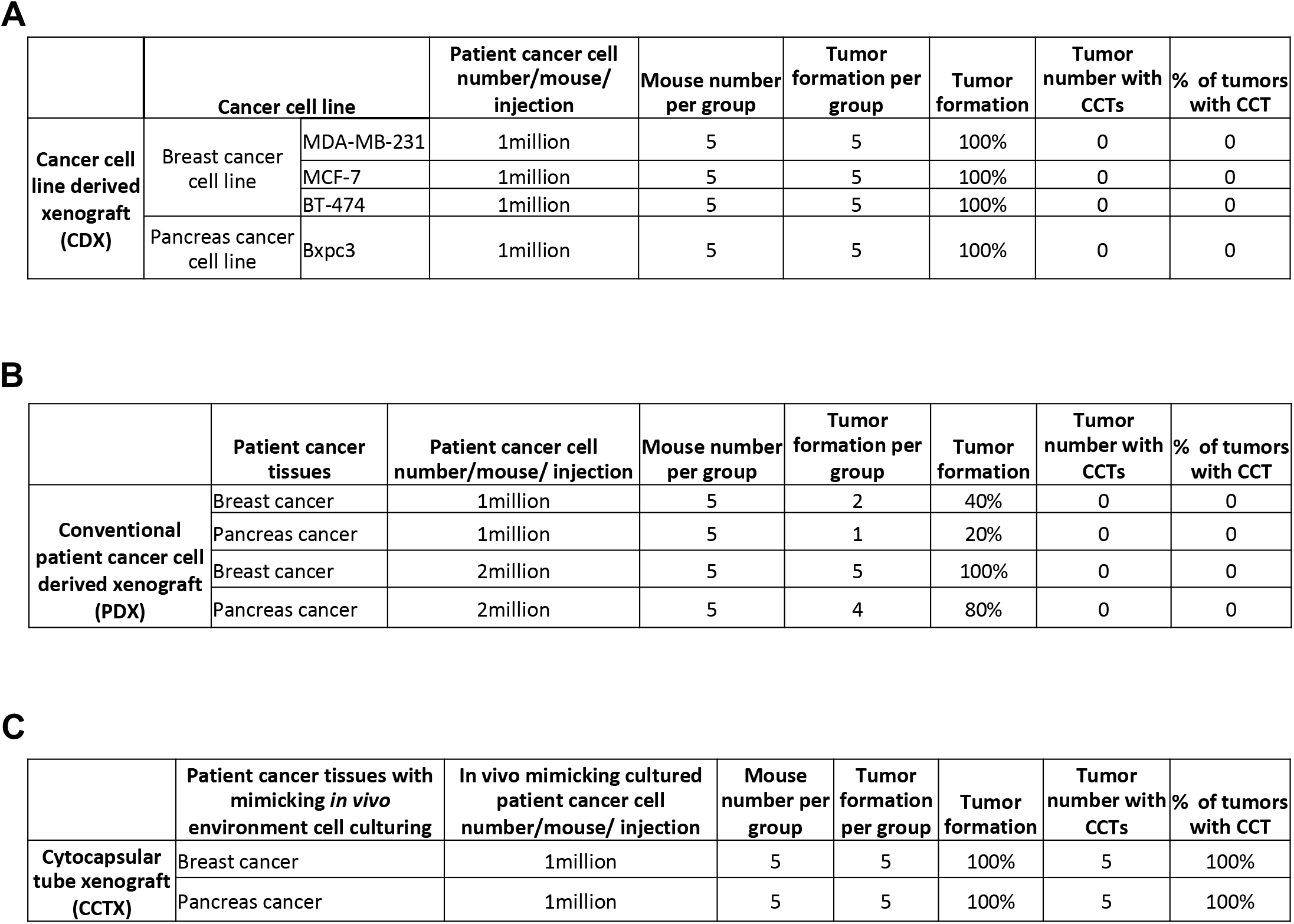
Analyses of cytocapsular tubes (CCTs) in conventional cancer cell line derived xenograft (CDX), patient cancer cell derived xenograft (PDX), and CCT xenograft (CCTX) in animal models in mice. (A) Table of CCT analyses of CDX in animal model in mice. (B) Table of CCT analyses of PDX in animal model in mice. (C) Table of CCT analyses of CCTX in animal model in mice.

**Fig.5.**
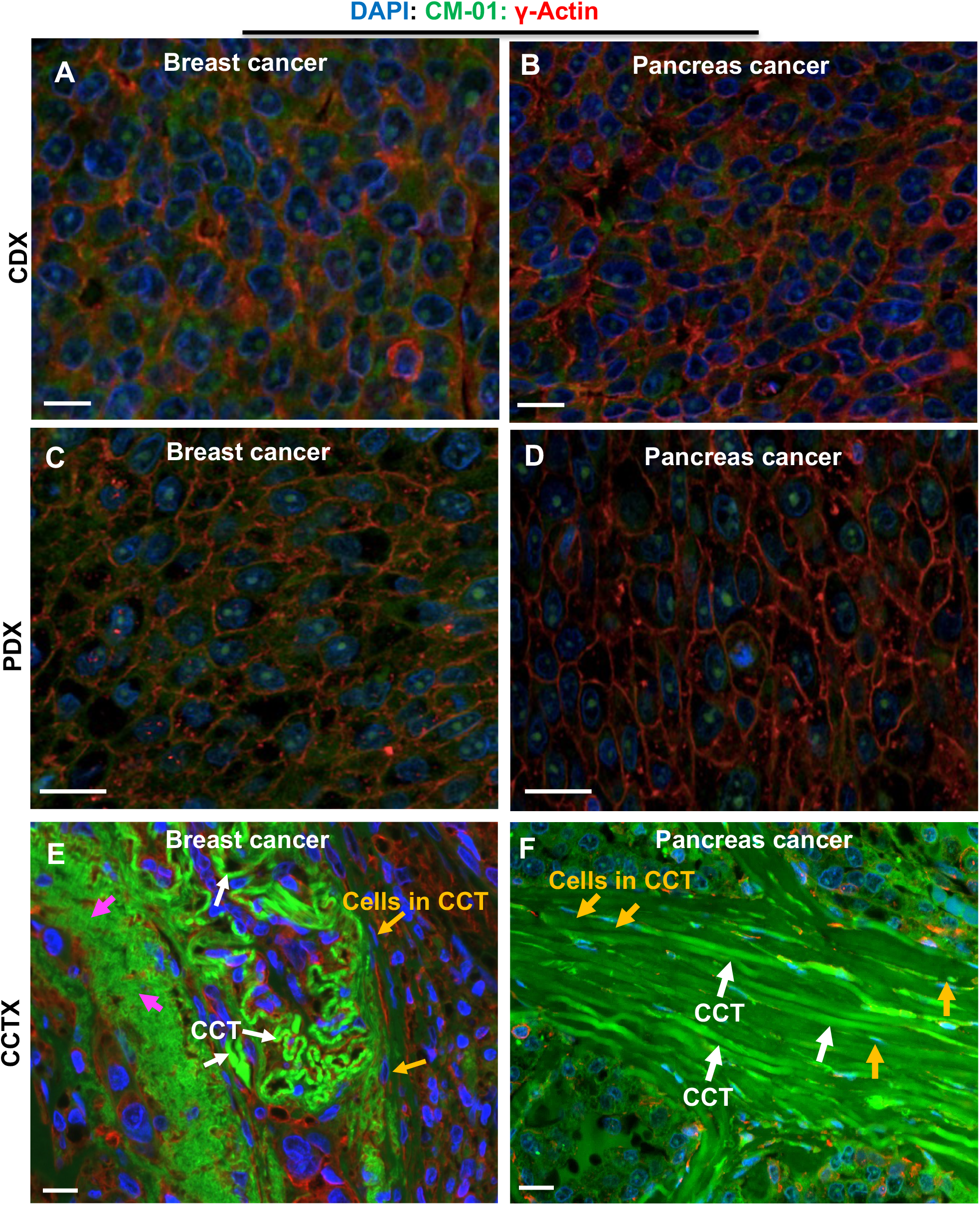
Analyses of cytocapsular tubes (CCTs) in conventional cancer cell line derived xenograft (CDX), patient cancer cell derived xenograft (PDX), and CCT xenograft (CCTX) with fluorescence microscope images. Representative fluorescence microscope image of FFPE sectioned tumor tissue slide of breast cancer and pancreas cancer from CDX (**A-B**), PDX (**C-D**) and CCTX (**E-F**) tumors from Tables A, B and C. There are a lot of compact cancer cells, but no CCTs in CDX (**A-B**) and PDX (**C-D**). (**E-F**) There are a lot of curved and winded and non-degraded cytocapsular tubes (CCTs, white arrows). Cancer cells in migration in CCTs (orange arrows) and many CCT strands (CCTS, purple arrows) are shown. The fluorescence channels of blue, green and red colors in images A-F were performed with the same exposure time, respectively. Scale bar, 10μm.

### Development of a cytocapsular tube xenograft (CCTX) platform

The understanding of the nature of cancer cell CCTs is currently in the very early stage and it is unknow how cancer cell CCTs are initiated, and it is also unknown what factors stimulate cancer cell CCT generation. We invented a series of conditions mimicking environments in tissues *in vivo* for cell culturing of cancer cells extracted from patient tumors, including: necessary growth factors, cell culture media, PH, and so on. Using fluorescence-activated cell sorting (FACS) assay with FITC-conjugated anti-CH-5 and PE-conjugated anti-CH-6 antibodies, we have isolated CH-5^high^/CH-6^high^ patient cancer cell subpopulations of breast cancer cells and pancreas cancer cells. In the xenograft assays (1million cancer cell/injection/mouse, 5mice/group), 100% of injections of CH-5^high^/CH-6^high^ patient cancer cell subpopulations of breast cancer cells and CH-5^high^/CH-6^high^ patient cancer cell subpopulations of pancreas cancer cells formed tumors. In these tumors, 100% of them harbor a lot of CCTs (**Fig. 4C, 5E-F**). The subsequent CCT analyses (**Table 1**) demonstrated that these xenografts with CH-5^high^/CH-6^high^ patient cancer cell subpopulations generate CCTs similar to those in clinical cancer tissues and were names CCTX for CCT xenograft. The further comprehensive comparisons of conventional cancer drug candidate development methods (2D and 3D *in vitro* methods, and CDX and PDX in animal models *in vivo*) and the newly developed cancer drug candidate development methods (CCT drug barrier screening platform *in vitro*, and CCTX animal model *in vivo*) suggest that the CCT drug barrier screening platform *in vitro* and CCTX animal model *in vivo* included method will be a highly efficient platform for cancer drug candidate development with high clinical prediction value (**Table 2**).

**Table 1.** Comprehensive comparison analyses of CCTs in the CCTX tumors and in the clinical breast cancer and pancreas cancer tissues.

|  |  | Clinical cancer tissue | CCT xenograft tissue |
| --- | --- | --- | --- |
| Breast cancer<br>(5 mice) | With cytocapsular tube (CCT) | Yes | Yes |
|  | CCT density (CCT/mm<sup>2</sup>) | 89-112 | 76-93 |
|  | CCT size in diameter/width (um) | 3-6 | 3-6 |
|  | Cancer cell migration in CCTs | Yes | Yes |
|  | with 3D CCT networks | Yes | Yes |
|  | With expression of CCT protein markers (CM-01, CM-02, CM-03) | 100% | 100% |
|  | CCT marker CM-01 abundance ratio to normal cells | 100 | 100 |
| Pancreas Cancer<br>(5 mice) | With cytocapsular tube (CCT) | Yes | Yes |
|  | CCT density (CCT/mm<sup>2</sup>) | 72-103 | 65-91 |
|  | CCT size in diameter/width (um) | 3-6 | 3-6 |
|  | Cancer cell migration in CCTs | Yes | Yes |
|  | with 3D CCT networks | Yes | Yes |
|  | With expression of CCT protein markers (CM-01, CM-02, CM-03) | 100% | 100% |
|  | CCT marker CM-01 abundance ratio to normal cells | 100 | 100 |

**Table 2.** Comprehensive comparison analyses of conventional cancer drug candidate development methods and highly efficient cancer drug candidate development methods. The last two line in the item 17 are blank as the clinical evaluations are still in the process and data are not available yet.

| Conventional cancer drug candidate development methods |  |  |  |  |  | Highly efficient cancer drug candidate development methods |  |  |
| --- | --- | --- | --- | --- | --- | --- | --- | --- |
|  |  | In vitro cancer cell line based |  | In vivo animal model |  | In vitro cancer cell line based | In vitro CCT drug barrier test | In vivo cytocapsular tube xenograft (CCTX) animal model |
|  | Important aspects of cancers in clinical cancer patients in vivo | 2D | 3D colony/organoid | CDX (Cell line derived xenografts) | PDX (patient tumor cell derived xenografts) | 2D | 3D CCT drug barrier screening platform | CCTX |
| 1 | With cytocapsular tube (CCT) drug barrier | No | No | No | No | No | Yes | Yes |
| 2 | With normal cancer cell in vivo microenvironments | No | No | No | No | No | Yes | Yes |
| 3 | With mimicked aneuploidy and genetic heterogeneity | Less | Less | Less | Less | Less | Yes | Yes |
| 4 | With cell proliferation | Yes | Yes | Yes | Unstable | Yes | Yes | Yes |
| 5 | With similar cell proliferation rate as in vivo | No | No | No | No | No | No | Yes |
| 6 | With cell viability | Yes | Yes | 100% | 0%-60% | Yes | Yes | 100% |
| 7 | With tumor formation success | - | - | 100% with 1million cells | 0-40% with 1million cells | - | - | 100% |
| 8 | with CCT in xenograft tumors | - | - | No | No | - | - | Yes |
| 9 | With CCT drug resistance | No | No | No | No | No | Yes | Yes |
| 10 | With CCT cancer metastasis pathways | No | No | No | No | No | Yes | Yes |
| 11 | Passed hits overcome CCT drug barriers in vivo | No | No | No | No | No | Yes | Yes |
| 12 | Passed hits overcome cancer metastasis in CCTs | No | No | No | No | No | Yes | Yes |
| 13 | Passed hits inhibit cancer cells in CCT | No | No | No | No | No | Yes | Yes |
| 14 | Partly mimic Pharmacokinetics and cytotoxicity | No | No | Yes | Yes | No | No | Yes |
| 15 | Partly mimic drug loss as in vivo | No | No | Yes | Yes | No | No | Yes |
| 16 | Test performance time | 1 week | 2 weeks | 3-5months | 6-10months | 1 week | 1 week | 4-8months |
| 17 | Clinical prediction percentage | <0.5% | <0.5% | 2%-3.5% | 3%-4% | <0.5% |  |  |

## Discussion

Cancer drug pan-resistant tumor metastasis (cdp-rtm) is a major source of cancer lethality^1–6^. Thus, we investigated CCT functions in cdp-rtm. Our studies demonstrated that superlarge membrane systems of cancer cell CCTs and its networks function as superdefence to conventional cancer drugs and conduct cdp-rtm that causes cancer lethality. We have developed a highly efficient CCTX for the development of precise and efficient cancer drugs.

### CCTs and all solid cancers are targetable and druggable

Obviously, cancer drugs can work only when they access the tumor cells. In clinical cancer tissues, CCTs and their networks are 3D in structure, superlarge and membrane-enclosed systems composed of superlarge, bi-layer phospholipid biomembranes. CCTs and networks provide with: 1) ultra-large, substantial and physical drug barriers protecting cancer cells inside, 2) membrane-enclosed 3D freeway systems reaching neighboring and far-distance destinations/organs, 3) cancer cells can migrate bi-directionally and proliferate in CCT lumens, 4) cancer cells can form new tumors in the large cytocapsulae that are connected to CCT networks, and the colony tumor cells can easily dissemble and leave the colonies and disperse to other sites via the CCT networks. CCTs and their networks in clinical tissues significantly increase drug resistance and challenges of cancer pharmacotherapy due to these aspects: 1) Physical drug barrier to cancer drugs that does not penetrate or eliminate CCTs: only if the drugs can penetrate the membrane systems of CCTs and networks, they can access and inhibit cancer cells in CCT lumens; 2) Separation of affected CCTs: when cancer drugs penetrate into parts of CCT fragments, they may lead to degradation of these CCT fragments. The unaffected CCT fragments can dynamically fuse and close the CCT fragment ends, and separate the unaffected CCTs from the affected CCT fragments. The unaffected CCTs and its networks continue to protect cancer cells inside; 3) Damage resilience of CCT networks: degradation in parts of CCT networks makes CCT cavities, but the survived CCTs still interconnect and form networks displaying CCT network damage resilience and cancer cell continue metastasis in the remaining CCT networks; 4) The heterogenicity of CCT densities and networks: CCT densities are highly various in clinical tissues, the higher CCT density, the lower is the average cancer drug dosage available to cancer cells, thus making these drugs less effective; 5) potent capacities of cancer cells to generate new CCTs; 6) The heterogenicity of CCT membrane proteins and lipids: CCTs from different organs, tissues and cells have heterogeneous membrane proteins (in protein types, species and numbers) and lipids (components and dynamic textures), and display quite various cancer drug barrier effects, and 7) acquired drug resistance of CCT membranes.

CCTs and networks display potent resistant capacities to many and even most of currently available drugs that are used commercially. Approximately 10million of cancer lethality occur after pharmacotherapies every year worldwide^1^. Here, we have successfully developed a CCT molecular marker protein CH-01 that is targeted by a small molecule CP-11, which shows potent capacities to successfully eliminate CCT barrier and kill cancer cells inside CCTs even with decreased dosage (by less than 33%). Our study evidenced that CCTs are targetable and druggable. It is expected that all solid cancers are druggable with precise and efficient cancer drugs aimed to eliminate CCTs and eradicate cancer cells in CCTs and their networks.

### Conventional low efficiency drug screening methods would need to be updated by including elements of CCTs and their networks

It is well-knowns that the current cancer drug development systems have very low (only average 3%) clinical prediction value^2–6^, but the underlying reasons are largely elusive. Besides the drug delivery pathway loss (physical, chemical and biological functional loss, average 93%-95% loss), CCTs and network drug barriers are a major source of functional loss of drugs that reach the cancer areas. Here, we showed that conventional CDX and PDX assays do not exhibit CCTs in animal models when using conventional cell culturing methods do not generate CCTs in these xenografts. This is consistent with the constantly very low clinical prediction value of the modern cancer drug screening and development methods. The absence of CCTs in conventional screening methods (2D cell culture and 3D cancer cell colonies and organoids *in vitro*, and CDX and PDX in animal models *in vivo*), require long lasting time research and development (10-12 years) and huge costs (up to $2.7billion/cancer drug), but obtain very low clinical prediction value, based on the existing animal models. Thus, it is necessary to include the CCT element in as more steps as possible in the screening of cancer drug candidates in order to increase efficiency of cancer drug discovery.

### CCTX is a highly efficient cancer drug development platform and CP-11 is a promising cancer drug candidate and booster drug candidate

The facts of: 1) cancer drug pan-resistant tumor metastasis (cdp-rtm) is a major source of cancer lethality and 2) CCT and networks are substantial, complex and dynamical membrane systems with pleiotropic functions^1-6^, which make CCT an indispensable component to overcome in order to effectively inhibit cancers. Here, we show the CCT drug barrier screening platform *in vitro* and the CCTX platform in animal model *in vivo*, and display potent inhibitory effects on cancer cells in CCTs. It is expected that drug candidate screening and discovery through CCT penetration assays *in vitro* and CCTX tests in animal models *in vivo*, which highly mimic clinical cancer tissue CCT situations and parts of the drug delivery pathways, will reach significantly increased or even high clinical prediction value. We understand that some conventional cancer drugs display excellent inhibitory effects on naked cancer cells without CCTs. It may be a desirable option to combine excellent conventional drugs with a booster cancer drug that specifically eliminates CCTs and their networks. Indeed, our studies (data will be reported in another manuscript) show that CP-11 displays potent abilities to function as booster drugs in combination with Sunitinib, Apalutamide, and Abiraterone acetate, and significantly boost these conventional cancer drugs inhibitory effects on cancer cells in CCTs.

In summary, our studies demonstrate that CCTs and networks function as superdefence against conventional cancer drugs and conduct conventional cancer drug pan-resistance tumor metastasis (cdp-rtm), CCTs are targetable and druggable, CCTX is a highly efficient cancer drug test platform, and the small molecule CP-11 is a promising cancer drug candidate and booster drug candidate.

## Methods and Materials

### Cells, antibodies, reagents and mice

Formalin-fixed paraffin-embedded (FFPE) clinical tissue slides were ordered from US Biomax. All cancer cell lines of MDA-MB-231, MCF-7, BT-474, LnCap and Bxpc3 and their specific cell culture medias as well as supplements were ordered from ATCC. Compounds of Sunitinib, Apalutamide, and Abiraterone acetate were ordered from Sigma. Small molecule CP-11 is self-synthesized. All compounds were dissolved in DMSO. Monoclonal mouse anti-γ-actin antibodies were ordered from Abcam (ab123034). Polyclonal and monoclonal rabbit anti-CM-01 (code, not real protein name) antibodies were self-developed. Polyclonal and monoclonal rabbit anti-CH-5 and anti-CH-6 (codes, not real protein names) antibodies were self-developed. Secondary antibodies of anti-mouse and anti-rabbit antibodies were ordered from Thermo Fisher Scientific. DAPI was ordered from VWR. Matrigel matrix was ordered from Sigma or Corning. *NOD-SCID* (strain name: *NOD.CB17-Prkdcscid/J*) and NCr-nu/nu mice were ordered from The Jackson Laboratory or Charles River Laboratories. All mouse work were reviewed and permitted by Cytocapsula Research Institute Animal Committee and performed in Charles River Laboratories.

### Immunohistochemistry (IHC) fluorescence staining assay

The sectioned FFPE clinical cancer tissue specimens were subjected to double immunohistochemistry staining with anti-CM-01 (1:200) and anti γ-actin (1:200) primary antibodies and DAPI (1:1,000) for 4h at 4 °C, followed by incubation with appropriate secondary antibodies for 1h at 4 °C in a dark room. Fluorescence images were taken with a Nikon 80i upright microscope with a 20× or 40× lens. All images were obtained using MetaMorph image acquisition software and were analyzed with ImageJ software.

### Cell viability assay

The pancreas cancer Bxpc3 cells and prostate cancer LnCap cells (5 × 10^3^) on 2D cell culture or 3D Matrigel matrix layers with cytocapsulae/cytocapsular tubes were treated with indicated compounds with series of concentrations for 72 h. Cell viability assays were performed on the cells with CellTiter-Glo luminescent cell viability assay kit (Promega) according to the manual. Three independent experiments were performed. Average IC_50_ results were shown (mean ± SD, *t*-test, two-tailed).

### Cell cytometry

Patient cancer cells (breast cancer cells and pancreas cancer cells) were extracted from the fresh clinical cancer tissues, and cultured with culturing situations mimicking *in vivo* environments: 37°C, 5% CO_2_, humidity and a series of other conditions. The CH-5^high^/CH-6^high^ patient cancer cell subpopulations were isolated by fluorescence-activated cell sorting (FACS) assay with self-developed FITC-conjugated anti-CH-5 and PE-conjugated anti-CH-6 antibodies.

### Tumor xenograft formation assay

In the tumor xenografted assay, the indicated cancer cell line cells, patient cancer cells, and CH-5^high^/CH-6^high^ subpopulation cancer cells (in the indicated cell numbers) were mixed with 100μl Matrigel/cell culture media mixture (Matrigel: cell culture media = 1:2) (Sigma or Corning). Indicated cancer cells/Matrigel/ cell culture media mixtures were injected into *NOD/SCID* female (for breast cancer xenograft) or NCr-nu/nu mice (for pancreas cancer xenograft) (the Jackson Laboratory) by subcutaneous injection. After the tumor formation (about 75 mm^3^ in volume, 5 mice/group), mice were sacrificed and tumors were excised. Tumor tissue samples were used for immunohistostaining and CCT analyses. The mouse experiments were performed according to the policies of Cytocapsula Research Institute Animal Committee and Charles River Laboratories.

### Statistical analysis

Quantitative data were statistically analyzed (mean ± SD, *t*-test, two-tailed). Statistical significance was determined by *t*-test. Significance was expressed as: *:*p* <0.1; **:*p* < 0.05; ***: *p* < 0.01, or with the *p*-value. *P* < 0.05 was considered significant.

Supported by a grant (to Dr. Yi) from Cytocapsula Research Institute Fund for Cytocapsular Tube Conducted Cancer Metastasis Research, and a grant (to Dr. Yi) from Chalst Inc Fund for Cytocapsular Tube Cancer Metastasis Research.

We greatly acknowledge Dr. Ed Harlow of Harvard Medical School (USA) and Cancer Institute of University of Cambridge (UK), and Dr. Nahum Sonenberg of McGill University of Canada, Dr Yubo Yang, and Dr. Qiping Hou for their help and meaningful discussion in the study.

## Notes

### Competing Interest Statement

The authors have declared no competing interest.

## References

1. Sung H, et al, Global cancer statistics 2020: GLOBOCAN estimates of incidence and mortality worldwide for 36 cancers in 185 countries. CA Cancer J Clin 2021; Online ahead of print.

2. Souza J, et al. Global health equity: cancer care outcome disparities in high-, middle-, and low-income countries. J Clin Oncol. 2016 Jan; 34(1): 6–13.

3. Chaffer CL, Weinberg RA. A Perspective on Cancer Cell Metastasis. Science 2011; 331:1559–64

4. Hanahan D, Weinberg RA. Hallmarks of cancer: the next generation. Cell 2011; 144:646–74.

5. Turajlic S, Swanton C. Metastasis as an evolutionary process. Science 2016; 352:169–75.

6. Chiang AC, Massagué J. Molecular basis of metastasis. N Engl J Med 2008;359: 2814–23.

7. Piet Borst. Cancer drug pan-resistance: pumps, cancer stem cells, quiescence, epithelial to mesenchymal transition, blocked cell death pathways, persisters or what? Open Biol. 2012 May; 2(5): 120066.

8. Harris AL. 1985 DNA repair and resistance to chemotherapy. Cancer Surv. 4, 601–624.

9. Kaelin Jr WG. 2005 The concept of synthetic lethality in the context of anticancer therapy. Nat. Rev. Cancer 5, 689–698.

10. Tuveson D, Hanahan D. 2011 Translational medicine: cancer lessons from mice to humans. Nature 471, 316–317.

11. Szakacs G, Paterson JK, Ludwig JA, Booth-Genthe C, Gottesman MM. 2006 Targeting multidrug resistance in cancer. Nat. Rev. Drug Discov. 5, 219–234.

12. Haimeur A, Conseil G, Deeley RG, Cole SP. 2004 The MRP-related and BCRP/ABCG2 multidrug resistance proteins: biology, substrate specificity and regulation. Curr. Drug Metab

13. Aas, T., Borresen, A. L., Geisler, S., Smith-Sorensen, B., Johnsen, H., Varhaug, J. E., et al. (1996). Specific P53 mutations are associated with de novo resistance to doxorubicin in breast cancer patients. Nat. Med. 2, 811–814.

14. Apperley, J. F. (2007). Part I: mechanisms of resistance to imatinib in chronic myeloid leukaemia. Lancet Oncol. 8, 1018–1029.

15. Gupta G and Massague J. Cancer metastasis: building a framework. Cell, 2006 Nov 17;127(4):679–95.

16. Yi F, Wagner G. Cytocapsular tubes conduct cell translocation. Proc Natl Acad Sci U S A 2018; 115: E1137–E1146.

17. Yi F, Wagner G, Cancer Metastases of 202 Kinds of Cancers via Cytocapsular Tubes, BioRxiv, 2021, https://doi.org/10.1101/2021.05.13.443996

18. Yi F, et al, Cytocapsular Tube-Based Precise Cancer Diagnosis in Colon Cancers. doi: https://doi.org/10.1101/2021.07.28.21261279.

19. Huerta S, et al, Colon cancer and apoptosis. Am J Surg 2006 Apr;191(4):517–26.

20. Mody K, Bekaii-Saab T. Clinical Trials and Progress in Metastatic Colon Cancer.

